# Population structure, genetic connectivity, and adaptation in the Olympia oyster (*Ostrea lurida*) along the west coast of North America

**DOI:** 10.1101/414623

**Authors:** Katherine Silliman

## Abstract

Effective management of threatened and exploited species requires an understanding of both the genetic connectivity among populations and local adaptation. The Olympia oyster (*Ostrea lurida*), patchily distributed from Baja California to the central coast of Canada, has a long history of population declines due to anthropogenic stressors. For such coastal marine species, population structure could follow a continuous isolation-by-distance model, contain regional blocks of genetic similarity separated by barriers to gene flow, or be consistent with a null model of no population structure. To distinguish between these hypotheses in *O. lurida*, 13,444 single-nucleotide polymorphisms (SNPs) were used to characterize rangewide population structure, genetic connectivity, and adaptive divergence. Samples were collected across the species range on the west coast of North America, from southern California to Vancouver Island. A conservative approach for detecting putative loci under selection identified 288 SNPs across 129 GBS loci, which were functionally annotated and analyzed separately from the remaining neutral loci. While strong population structure was observed on a regional scale in both neutral and outlier markers, neutral markers had greater power to detect fine-scale structure. Geographic regions of reduced gene flow aligned with known marine biogeographic barriers, such as Cape Mendocino, Monterey Bay, and the currents around Cape Flattery. The outlier loci identified as under putative selection included genes involved in developmental regulation, sensory information processing, energy metabolism, immune response, and muscle contraction. These loci are excellent candidates for future research and may provide targets for genetic monitoring programs. Beyond specific applications for restoration and management of the Olympia oyster, this study lends to the growing body of evidence for both population structure and adaptive differentiation across a range of marine species exhibiting the potential for panmixia. Computational notebooks are available to facilitate reproducibility and future open-sourced research on the population structure of *O. lurida*.

## INTRODUCTION

Coastal marine ecosystems provide important services such as carbon sequestration, food production, and recreation (Luisetti et al., 2014), yet contain some of the most exploited and threatened species on earth. As evidence for the direct impacts of human activities (e.g., over-harvesting, increasing atmospheric CO_2_, and nutrient run-off) on these species grows, there has been increased focus on restoring depleted abundances, recovering ecosystem services, and determining which species are capable of adapting to environmental change (Granek et al., 2010). Effective management of threatened and exploited species requires an understanding of both the genetic connectivity among populations and adaptation across environmental gradients (Baums, 2008; Miller and Ayre, 2008; Palumbi, 2003). Anthropogenic movement of individuals between populations, either for aquaculture or restoration purposes, can confound signals of population structure and should be evaluated when drawing conclusions about genetic connectivity (David, 2018). For the numerous coastal marine species with planktonic dispersal, high connectivity can result in subtle among populations can obscure population boundaries and oppose the diversifying effects of natural selection through gene flow (Lenormand, 2002). A growing body of evidence indicates that both limited effective dispersal and local adaptation may be more common in marine species than previously hypothesized (Cowen et al., 2000; Hauser and Carvalho, 2008; Sanford and Kelly, 2011; Weersing and Toonen, 2009).

Neutral molecular markers (e.g., microsatellites) have traditionally been used exclusively to identify the geographic structure of subpopulations and estimate the genetic connectivity between them (Funk et al., 2012); however, these do not give insight into the scale or magnitude of adaptive divergence. For populations connected by dispersal, adaptation to local conditions can still occur if the strength of selection overcomes the homogenizing effect of gene flow (Hellberg, 2009). Advances in genomic and computational techniques, such as genotype-by-sequencing (GBS), have facilitated the detection of genomic regions that may be influenced by natural selection (“outlier loci”) (Stapley et al., 2010). Although often referred to as adaptive markers, these outliers may only be linked to loci that are under natural selection rather than confer an adaptive advantage themselves. Outlier loci have provided increased spatial resolution of population structure compared to neutral loci alone for some marine species (Drinan et al., 2018; Van Wyngaarden et al., 2016; Milano et al., 2014), but not all (Moore et al., 2014). In addition to potentially resolving fine-scale genetic differentiation, outlier loci may be useful for characterizing the adaptive potential of a species or population to future environmental conditions (Eizaguirre and Baltazar-Soares, 2014). Inferences from both neutral and adaptive markers should be combined when making management recommendations (Funk et al., 2012).

The Olympia oyster (*Ostrea lurida*, Carpenter 1864) is a native estuarine bivalve found from Baja California to the central coast of Canada, patchily distributed over strong environmental gradients (Chan et al., 2017; Schoch et al., 2006). Oysters are ecosystem engineers in estuaries, providing structured habitat and removing suspended sediments (zu Ermgassen et al., 2013; Coen et al., 2011). Unlike other oysters where both males and females spawn gametes (e.g., *Crassostrea*), the females fertilize eggs with sperm from the water column and initially brood larvae in the mantle cavity. After release, the larvae have been reported to be planktonic from seven days to eight weeks before settling on a hard substrate (Baker, 1995). The impact of maternal brooding on population structure in Osterideae has not been examined.

Following devastating commercial exploitation in the 19^th^ and early 20^th^ centuries, recovery of Olympia oyster populations has been stifled by other anthropogenic threats (e.g., water quality issues, habitat loss, and possibly ocean acidification (Blake and Bradbury, 2012; Hettinger et al., 2013; Sanford et al., 2014)). The last 15 years has seen increased interest in the Olympia oyster, with restoration projects underway by both government and nongovernment agencies across its range (Pritchard et al., 2015). Current knowledge about the population genetic structure of *O. lurida* comes primarily from an unpublished 2011 dissertation, which sampled from San Francisco, CA to Vancouver Island, BC and found regional population structure using microsatellites (Stick, 2011). Two phylogeographic studies using two mitochondrial loci identified a phylogeographic break north of Willapa Bay, WA and established the southern boundary divide between *O. lurida* and its sister species *Ostrea conchaphila* (Polson et al., 2009; Raith et al., 2016). Future and ongoing management plans would benefit greatly from thorough analysis of the fine-scale genetic structure using modern genomic techniques and rangewide sampling (Camara and Vadopalas, 2009).

The objective of this study was to characterize the spatial population structure of the Olympia oyster across the majority of its range using both neutral and adaptive markers derived from genome-wide single nucleotide polymorphisms (SNPs). I specifically tested whether patterns of genetic variation suggest a smooth continuum of allele frequency shifts consistent with isolation-by-distance (IBD) (Malécot, 1968), regional blocks of genetic similarity that correspond to physical barriers (Hare and Avise, 1996), or the null model of no significant genetic differentiation (Grosberg and Cunningham, 2001). SNPs produced from high-throughput sequencing have led to the identification of previously undetected population structure in a number of marine and terrestrial species (Everett et al., 2016; Reitzel et al., 2013; Van Wyngaarden et al., 2016). Compared to the Atlantic coast of North America (Hoey and Pinsky, 2018), studies utilizing genome-wide SNPs for marine taxa from the Pacific coast are far fewer in number and have been limited to regional spatial scales (De Wit and Palumbi, 2013; Drinan et al., 2018; Gleason and Burton, 2016; Larson et al., 2014; Martinez et al., 2017) or in the number of sampling sites (Pespeni et al., 2010; Tepolt and Palumbi, 2015). This study is the first of my knowledge to utilize thousands of SNPs to extensively survey the rangewide population structure of a marine species along this coast.

A secondary aim of this study was to produce a reproducible computational pipeline to go from raw data to results and figures, using Jupyter notebooks. Jupyter notebooks are interactive documents that integrate text, code, and analysis results (Kluyver et al., 2016). A major issue for genomic analyses today is how to clearly explain the computational methods used in order to allow for reproducibility (Kanwal et al., 2017). This open access pipeline is intended to provide an example template to improve reproducibility in future studies and function as an instructional tool for biologists and early career scientists who wish to apply these methods to their own study organisms.

## MATERIALS AND METHODS

### Sample collection

20-25 adult *Ostrea lurida* over 2cm in length were collected primarily by hand from the intertidal (approx. 0m to −1m tidal height) at 20 sites ranging from Klaskino Inlet, Vancouver Island (50° 17’ 55”) to San Diego Bay, CA (32°361’ 9”) in 2014 (Figure 1, Table A1). When possible, oysters were sampled randomly along 10m transects. Each site represents a separate bay, except for Willapa Bay, WA and San Francisco Bay, CA which had two sampling sites each. In the case of one site from Willapa Bay, WA, oysters were collected subtidally (approx. 8m depth) through dredge harvesting of the Pacific oyster, *Crassostrea gigas*. Adductor tissue samples were preserved in RNALater, followed by storage in −80°C. DNA was isolated using DNeasy Blood & Tissue kits (Qiagen) and E.Z.N.A. Mollusc DNA kits (Omega Bio-Tek) with RNAse-A treatment following manufacturer instructions and limiting tissue digestion time to no more than 90 minutes. All DNA samples were quantified using the Qubit dsDNA BR Assay kit (Life Technologies) on a Qubit v2.0 (Life Technologies). DNA quality was verified by agarose gel electrophoresis of 1-2 uL extracted DNA on an 0.8% TAE gel.

**Figure 1:**
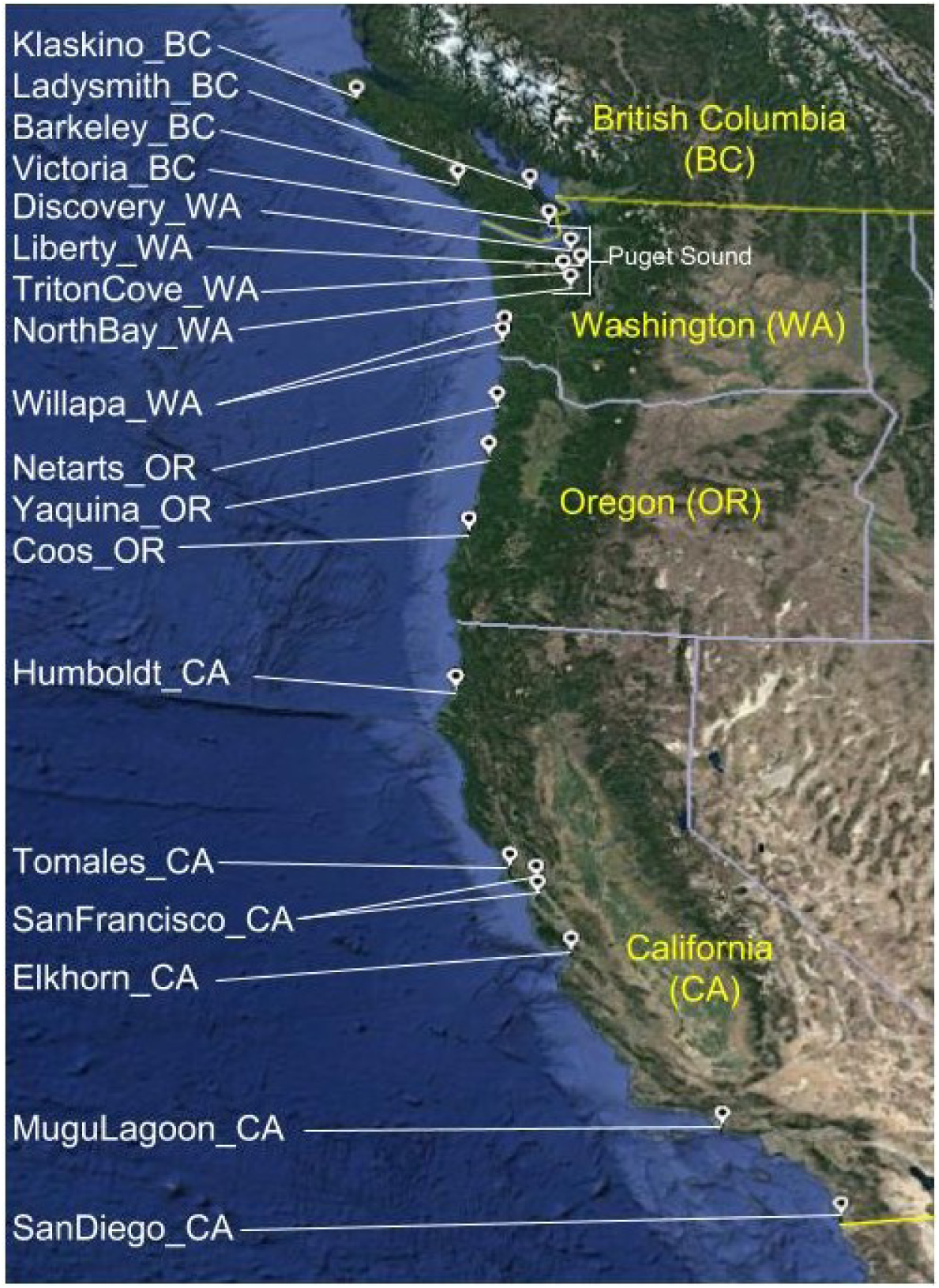
Map of 20 Olympia oyster (*Ostrea lurida*) collection sites from the west coast of North America.

### Genotype-by-sequencing analysis

Library preparation for genotype-by-sequencing (GBS) followed the protocol by Elshire et al. (2011) using the *ApeKI* restriction enzyme, with an additional size selection step and slight modifications to PCR amplification. A detailed protocol can be found at [redacted]. All libraries were size-selected for fragment sizes between 200bp-450bp on a Blue PippinPrep (Sage Science) to reduce the number of loci sequenced and ensure adequate sequencing coverage. One pool of 90 samples was sequenced across two 100bp paired-end Illumina HiSeq 2500 lanes, with only the forward sequencing read used for analysis. Seven other pools with a maximum of 48 libraries each were sequenced on eight 100bp single-end lanes (246 different samples, 86 technical replicates, 332 libraries total). Sampling sites were spread out across all libraries in order to minimize batch effects from library preparation and sequencing. Raw sequencing reads were de-multiplexed, quality filtered, and de novo clustered using the API version of the seven step computational pipeline *ipyrad* v.0.7.24 (Eaton, 2014) and implemented in Python via a Jupyter Notebook running on a large computational cluster. Demultiplexing (Step 1) used sample-specific barcode sequences, allowing one mismatch in the barcode sequence. Base calls with a Phred quality score under 20 were converted to Ns, and reads containing more than 5 Ns were discarded. Adapter sequences, barcodes, and the cutsite sequences were trimmed from filtered reads, with only reads greater than 35bp retained (Step 2). Reads were then clustered using a sequence similarity threshold of 85% both within (Step 3) and between samples to genotype polymorphisms (Steps 4, 5) and identify orthologous loci (Step 6) with a minimum of 10x read coverage. Replicate samples were assembled separately and then compared using custom Perl scripts by Mikhail Matz (Wang et al., 2012). The replicate with the largest number of GBS loci after final filtering (Step 7) was retained. Samples were removed if they had fewer than 200,000 raw sequencing reads, fewer than 15,000 assembled clusters of at least 10x read depth, and were missing data for over 55% of loci assembled across at least 75% of samples, with Steps 4-7 rerun using the remaining individuals.

The final assembly was then filtered for excess heterozygosity based on deviations from Hardy-Weinberg equilibrium (HWE) in at least two populations, sample coverage of 75%, and an overall minor allele frequency (MAF) of 2.5%, retaining only GBS loci found in at least one individual from all populations. Preliminary analyses conducted on datasets allowing more or less missing data showed that the inferred population structure was robust to missing data up to 40%. Population genetic summary statistics, with the exception of *F*_*ST*_, did change quantitatively due to missing data but not qualitatively (not shown) (Cariou et al., 2016). Filtering steps were conducted using VCFtools (Danecek et al., 2011), custom Python code, and code adapted from Jon Puritz’s lab (Puritz et al., 2014). Input files and formats for subsequent analysis of population structure were created using a combination of custom Python code, custom R code, and the *radiator* R package (Gosselin, 2017). Every step of the assembly, filtering process, and creation of input files can be reproduced through Jupyter notebooks.

### Detection of loci under putative selection

Following recommendations to utilize multiple methods to detect loci under putative directional selection (Benestan et al., 2016; Rellstab et al., 2015), three approaches were used on the filtered SNP dataset: BayeScan v.2.1, *OutFLANK* v.0.2, and *pcadapt* v.4.0.2. For BayeScan and *OutFLANK*, individuals were grouped into populations by sampling site. GBS loci which had SNPs identified as outliers in at least two of the approaches were classified as putative adaptive GBS loci. From these GBS loci, any SNP that had been identified as an outlier by at least one approach was separated from the full SNP dataset to create an “outlier” SNP dataset. Subsequent analyses of population structure were conducted on three SNP datasets: all SNPs (combined), outlier SNPs, and neutral SNPs—which excluded any SNP found on a putative adaptive GBS locus.

BayeScan uses a Bayesian approach to apply linear regression to decompose *F*_*ST*_ coefficients into population-and locus-specific components and estimates the posterior probability of a locus showing deviation from Hardy–Weinberg proportions (Foll and Gaggiotti, 2008). BayeScan analysis was based on 1:100 prior odds, with 100,000 iterations, a burn-in length of 50,000, a false discovery rate (FDR) of 10%, and default parameters. Results were visualized in R. *OutFLANK* is an R package that identifies F_ST_ outliers by inferring a distribution of neutral *F*_*ST*_ using likelihood on a trimmed distribution of *F*_*ST*_ values. Because of its likelihood method, *OutFLANK* calculates *F*_*ST*_ without sample size correction when inferring the neutral distribution. Simulation studies have shown that this approach has lower false positive rates compared to other *F*_*ST*_ outlier methods (Whitlock and Lotterhos, 2015). *OutFLANK* was run using default parameters and a *q*-value threshold of 0.1, which can be considered a false discovery rate (FDR) of 10%. For the R package *pcadapt*, individuals are not sorted into predefined populations. Instead, *pcadapt* ascertains population structure using principal component analysis (PCA), then identifies markers under putative selection as those that are excessively correlated with population structure. When compared to BayeScan, *pcadapt* was shown to have greater power in the presence of admixed individuals and when population structure is continuous (Luu et al., 2017)—both scenarios which are likely in *O. lurida*. A scree plot representing the percentage of variance explained by each PC was used to choose the number of principal components (*K*) for *pcadapt*, and SNPs with a *q*-value threshold of 0.1 were categorized as outliers.

Putative adaptive GBS loci were functionally annotated through Blast2GO. Sequences were compared against molluscan sequences in GenBank’s *nr* database using the BLASTx algorithm with default parameters and a *e*-value hit filter of 10^-3^, and against EMBL-EBI InterPro signatures. Gene ontology terms were mapped to annotations with default parameters except for an *e*-value hit filter of 10^-3^ (Götz et al., 2008). Minor allele frequency was plotted against latitude individually for each outlier SNP in order to identify clinal patterns of allele frequency shifts.

### Summary statistics, population differentiation, and spatial structure

Population genetic summary statistics were calculated on the combined, neutral, and outlier datasets to describe and compare overall and population-specific genetic diversity. Observed heterozygosity (*H*_*o*_), expected heterozygosity (*H*_*e*_), overall *F*_*ST*_, and *F*_*IS*_ were calculated using the *basic.stats* function in the R package *hierfstat* (Goudet and Jombart, 2015). Confidence intervals for population-specific *F*_*IS*_ were determined using the *boot.ppfis* function in *hierfstat* with 1,000 bootstrap replicates. Pairwise *F*_*ST*_ following Weir and Cockerham (1984) was calculated using the *genet.dist* function in *hierfstat*. Heatmaps of pairwise *F*_*ST*_ values were created using *ggplot2* (Wickham, 2016). A Mantel test of coastal water distance (calculated by drawing routes between all sites on Google Earth) and *F*_*ST*_ */*1 *-F*_*ST*_ as implemented in *adegenet* tested for evidence of isolation-by-distance (Sokal, 1979).

Rangewide population structure of *O. lurida* was characterized using a combination of Bayesian clustering and multivariate ordination approaches. These methods were applied to both the outlier and neutral datasets. The model-based Bayesian clustering method STRUCTURE v.2.2.4 (Pickrell and Pritchard, 2012) as implemented in the *ipyrad* API was used to determine the number of distinct genetic clusters (*K*) with a burn-in period of 50,000 repetitions followed by 200,000 repetitions. Five replicate analyses were performed for each dataset with values of *K* = 1-10, with each replicate independently subsampling one SNP per GBS locus and using a different random seed. Replicates were summarized and visualized using the CLUMPAK server (Kopelman et al., 2015). The Δ*K* method implemented in STRUCTURE HARVESTER was used to determine an optimal *K* (Earl, Dent A. and vonHoldt, Bridgett M, 2012). PCA was implemented in the R package *adegenet* (Jombart and Ahmed, 2011) using “unlinked” datasets, where a single SNP with the least missing data across samples was chosen for each GBS locus (or the first SNP in the locus in the case of a tie). Missing data was filled by randomly drawing an allele based on the overall allele frequency across all individuals. The R package *PCAviz* was used to visualize PCA results and correlate PC loadings with latitude (Novembre et al., 2018). Results from STRUCTURE, PCA, and pairwise *F*_*ST*_ were used to identify phylogeographic “regions”. Summary statistics, including *H*_*o*_, *H*_*e*_, *F*_*IS*_, and *F*_*ST*_ were calculated for each region using using the *basic.stats* function in the R package *hierfstat* (Goudet and Jombart, 2015).

### Estimating connectivity and historical relationships

Spatial variation in gene flow and genetic diversity was calculated and visualized using the program EEMS (Estimated Effective Migration Surfaces) (Petkova et al., 2016). This method identifies geographic regions where genetic similarity decays more quickly than expected under isolation-by-distance based on sampling localities and a pairwise genetic dissimilarity matrix derived from SNP data. These regions may be interpreted as having reduced gene flow. A dissimilarity matrix was calculated for the neutral dataset using a variant of the *bed2diffs* R code included in the EEMS package that takes input from a *genind* R object. An outer coordinate file for defining the potential habitat of *O. lurida* was produced using the polyline method in the Google Maps API v3 tool (http://www.birdtheme.org/useful/v3tool.html). The habitat shape followed the shape of the coastline and excluded land regions that *O. lurida* larvae would not naturally be able to cross (e.g., the Olympic peninsula separating outer coast populations and those in Puget Sound, WA). The EEMS model is recommended to be run for various numbers of demes, which establishes the geographic grid size and resulting potential migration routes. Three independent analyses were run for each deme size (200, 250, 300, 350, 400, 500, 600, 700) for a total of 24 runs, with a burn-in of 1,000,000 and MCMC length of 5,000,000 iterations. The convergence of runs was visually assessed and results were combined across all analyses and visualized using the *Reemsplots* R package—producing maps of the effective diversity (*q*) and effective migration rate (*m*).

To infer the evolutionary relationship among sampling sites, including population splits and migration events, I reconstructed population graph trees using the software TreeMix (Pickrell and Pritchard, 2012). This method uses the covariance of allele frequencies between populations to build a maximum likelihood graph relating populations to their common ancestor, taking admixture events (”migration”) into account to improve the fit to the inferred tree. The population graph was rooted with the two southernmost *O. lurida* population (San Diego, CA and Mugu Lagoon, CA), then run allowing between 0-10 migration events. For each value of migration events, I calculated the proportion of variance in relatedness between populations that is explained by the population graph to evaluate model fit (Wang, 2017).

## RESULTS

### GBS and outlier detection

Almost all replicate pairs showed > 88% similarity in heterozygous calls except for one, in which case both replicates were dropped. 117 samples remained after removal of 14 samples with < 200,000 raw sequencing reads, 49 samples with < 15,000 clusters, and 65 samples missing data for over 55% of loci assembled across at least 75% of samples. One of the sampling sites for Willapa Bay, WA had a low number of individuals after filtering, so individuals from these two sites were combined into one population, for 19 total populations (4–9 individuals per population, mean = 6.2). 42,081 biallelic SNPs across 9,836 GBS loci were genotyped in greater than 75% of these individuals (2.8% of prefiltered loci assembled by *ipyrad*). Average read depth per individual per GBS locus ranged from 21 to 120 (mean = 32±14). Further filtering by HWE and MAF > 2.5% reduced the dataset to 13,444 SNPs across 6,192 GBS loci (the “combined” dataset).

Three different methods were employed to identify putative SNPs under selection. The number of outliers detected by each program and the overlap between programs is illustrated in Figure D1. *OutFLANK*, as the most conservative of the programs used (Whitlock and Lotterhos, 2015), had the lowest number of outlier markers detected with 31 SNPs across 16 GBS loci. 29 SNPs found across 15 GBS loci were identified as outliers by all three programs. 129 97 GBS loci contained SNPs identified as outliers by at least two approaches, with 288 168 SNPs included in the outlier dataset for subsequent population structure analyses. The neutral dataset, with 13,093 SNPs across 6,063 GBS loci, excluded any SNP found on a GBS locus with an outlier SNP.

### Summary statistics, population differentiation, and spatial structure

#### Summary statistics

Global *F*_*ST*_ for outliers (*F*_*ST*_ = 0.399) was almost four five times greater than for the combined and neutral SNPs (*F*_*ST*_ = 0.105 (combined), 0.097 (neutral)). The outlier dataset had the lowest *H*_*o*_, but the highest *H*_*e*_ (Table 1). Average *F*_*IS*_ within populations for the combined dataset was 0.0424, with all populations having a significantly positive *F*_*IS*_ value except Ladysmith, BC, Tomales Bay, CA, and South San Francisco Bay, CA which had small, yet significantly negative *F*_*IS*_ values. Mugu Lagoon had the highest *F*_*IS*_ value (Table A1). Summary statistics for the six phylogeographic regions identified in the following section are show in Table B1. Summary statistics were quantitatively very similar for the combined and neutral datasets, so that only the results for the outlier and neutral datasets are reported for all subsequent analyses.

**Table 1.** Overall summary statistics for the combined (13,444 SNPs), neutral (13,093 SNPs), and outlier (288 SNPs) datasets. *H*_*o*_, observed heterozygosity averaged across loci; *H*_*e*_, expected heterozygosity averaged across loci; *F*_*IS*_ & *F*_*ST*_, Wright’s *F*-statistics averaged across loci (Nei and Chesser, 1983); pairwise *F*_*ST*_, average of all pairwise *F*_*ST*_ values (Weir and Cockerham, 1984)

| Dataset | $H_o$ | $H_e$ | $F_{IS}$ | $F_{ST}$ | Avg. pairwise $F_{ST}$ | Avg. within-population $F_{IS}$<br>(Min - Max) |
| --- | --- | --- | --- | --- | --- | --- |
| Combined | 0.191 | 0.225 | 0.051 | 0.105 | 0.1026<br>($1.83e^{-3}$ - 0.1902) | 0.0424<br>(-0.0960 - 0.1327) |
| Neutral | 0.192 | 0.224 | 0.052 | 0.097 | 0.095<br>($1.38e^{-3}$ - 0.177) | 0.0425<br>(-0.0953 - 0.1288) |
| Outlier | 0.162 | 0.280 | 0.039 | 0.399 | 0.358<br>(0.0106 - 0.6030) | 0.0241<br>(-0.2649 - 0.3149) |

**Table 2.**
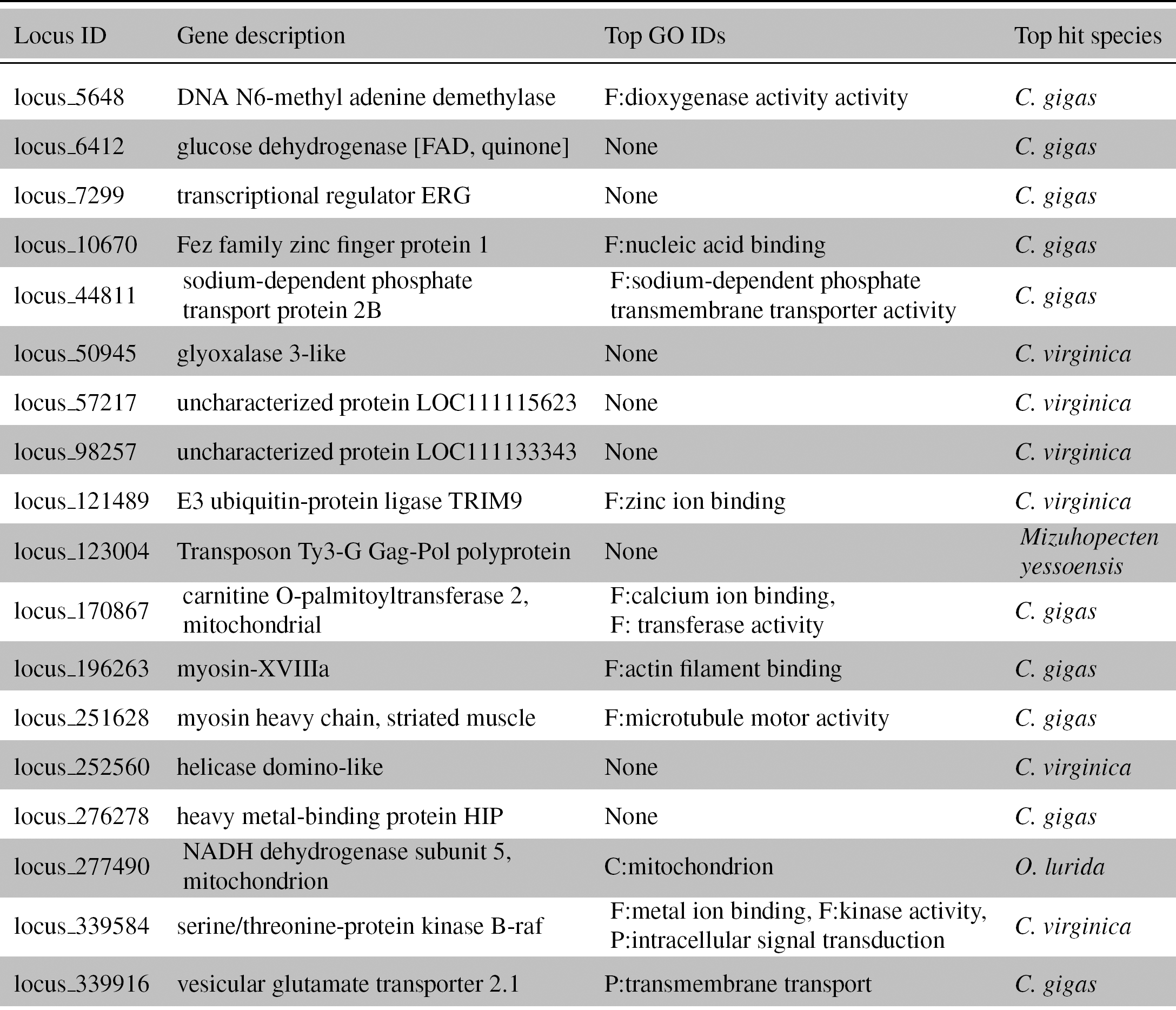
BLASTx and gene ontology (GO) annotation results for outlier loci. Only the 1814 loci with positive BLAST hits are shown. F: molecular function, C: cellular component, P: biological process.

#### Spatial structure

Bayesian population structure analysis from STRUCTURE differed slightly between the neutral and outlier datasets. For neutral SNPs, *K* = 5 had the strongest support based on the Evanno method. Visual inspection of STRUCTURE admixture plots for *K* = 6 included an additional population grouping of Willapa Bay, WA and Coos Bay, WA that was further supported by PCA and *F*_*ST*_ results (Figure 2). I refer to these six groupings as phylogeographic “regions”: Northwest Vancouver Island, BC (*NWBC*), Puget Sound, WA plus Victoria, BC and Ladysmith Harbour, BC (*Puget+BC*), Willapa Bay, WA plus Coos Bay, OR (*Willapa*), the other two Oregon sites (*Oregon*), Northern California (*NoCal*) from Humboldt Bay, CA to San Francisco Bay, CA, and Southern California (*SoCal*) from Elkhorn Slough, CA to San Diego Bay, CA. STRUCTURE results for the outlier SNPs supported *K* = 2, but visually results were similar between the outlier and neutral SNPs at *K* = 5 with the exception of Discovery Bay, WA in Puget Sound showing higher admixture with *NWBC* (Figure 2). The separation of *Willapa* sites from *Oregon* was not observed using the outlier dataset until *K* = 8.

**Figure 2:**
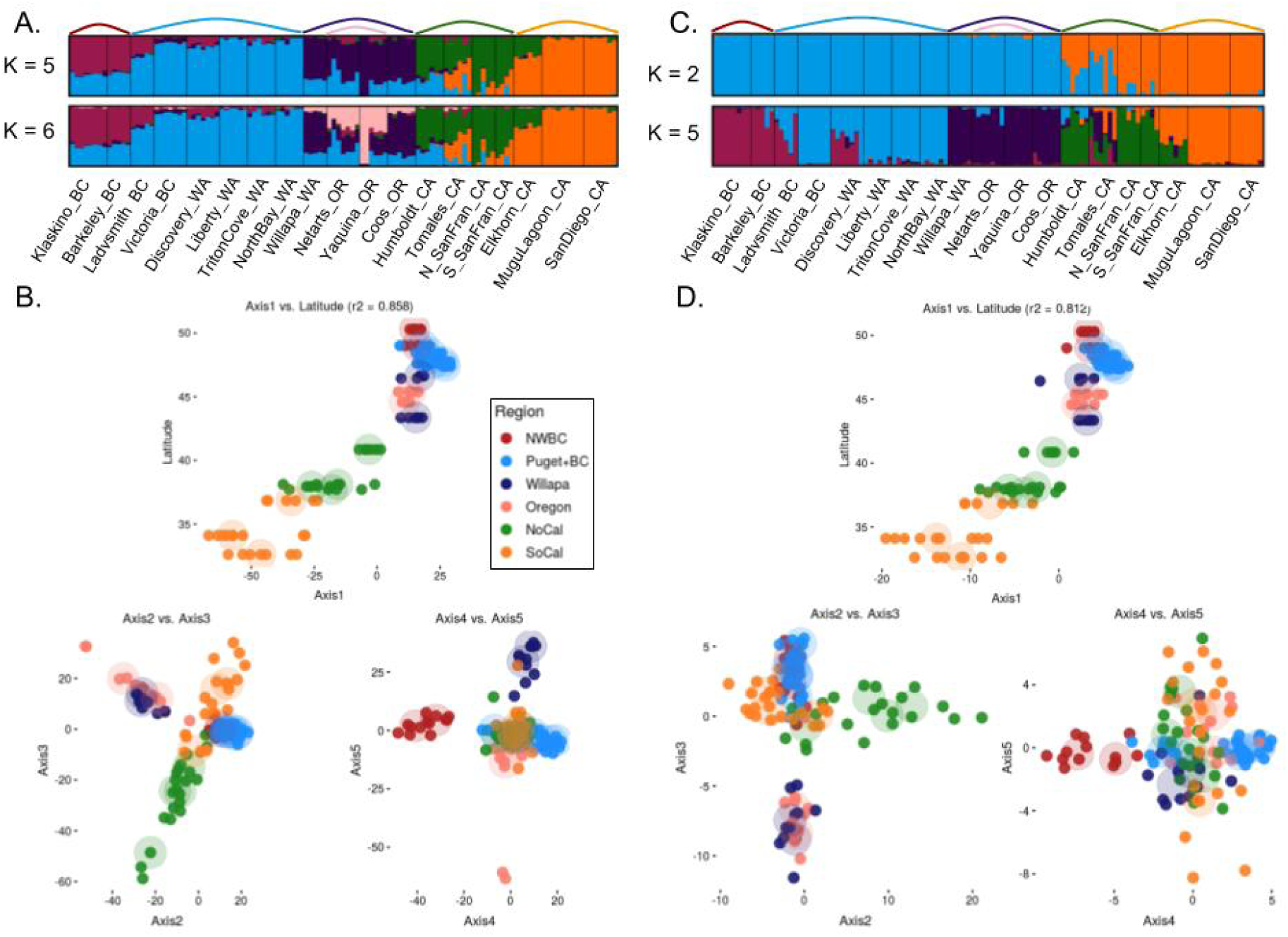
Population structure results for 19 *O. lurida* populations using (A-B) neutral loci and (C-D) outlier loci. (A, C) Plots of individual admixture determined using the program STRUCTURE at the *K* recommended by the ΔK method (*K* = 5 neutral, *K* = 2 outlier), as well as at the value of *K* inferred from PCA (*K* = 6 neutral, *K* = 5 outlier). (B, D) Principal components analysis plots for PCs 1-5. PC1 is plotted against latitude of sampling site, then PC2 vs. PC3 and PC4 vs. PC5. Large transparent circles indicate the centroid of populations. Colors refer to the phylogeographic regions of each population.

PCA on both SNP datasets demonstrated a strong relationship between latitude and the first principal component (PC) (neutral: *R*^2^ = 0.858, outlier: *R*^2^ = 0.806) (Figure 2). PCs 2-5 in the neutral dataset separated out individuals by phylogeographic region, with PC2 separating (*Puget+BC, NWBC*) and (*Willapa, Oregon*), PC3 separating *NoCal* and *SoCal*, PC4 separating *NWBC* and *Puget+BC*, and PC5 separating *Oregon* from *Willapa* (Figure 2). PC1 of the neutral dataset represented 5.8% of the total variance and PCs 2-5 represented 2.5%-1.5% of the variance. The outlier dataset showed similar regional spatial structure for PCs 2-4, but only showed slight separation of *Oregon* from *Willapa* (Figure 2). PC1 of the outlier dataset represented 21.4% of the total variance, and PCs 2-5 represented 9.2%-3.1% of the total variance.

#### Population differentiation and isolation-by-distance

Pairwise population-specific *F*_*ST*_ was higher for outlier SNPs, but both datasets qualitatively illustrated roughly six geographic “regions” of genetically similar populations and an overall trend of isolation-by-distance, where *F*_*ST*_ values were higher between sites that were farther away from each other (Figure 3, Tables C1 & C2). The three comparisons with the lowest pairwise *F*_*ST*_ using the neutral dataset were Willapa Bay, WA/Coos Bay, OR (0.0014), Mugu Lagoon, CA/San Diego Bay, CA (0.0016), and North/South San Francisco Bay, CA (0.0039). Victoria, BC showed higher pairwise *F*_*ST*_ with the other British Columbia sites than with sites from Puget Sound, WA. Mantel tests showed a significant correlation between pairwise *F*_*ST*_ and coastal water distance for both datasets, indicating a strong trend of isolation-by-distance (*p*-value = 0.001) (Figure 3).

**Figure 3:**
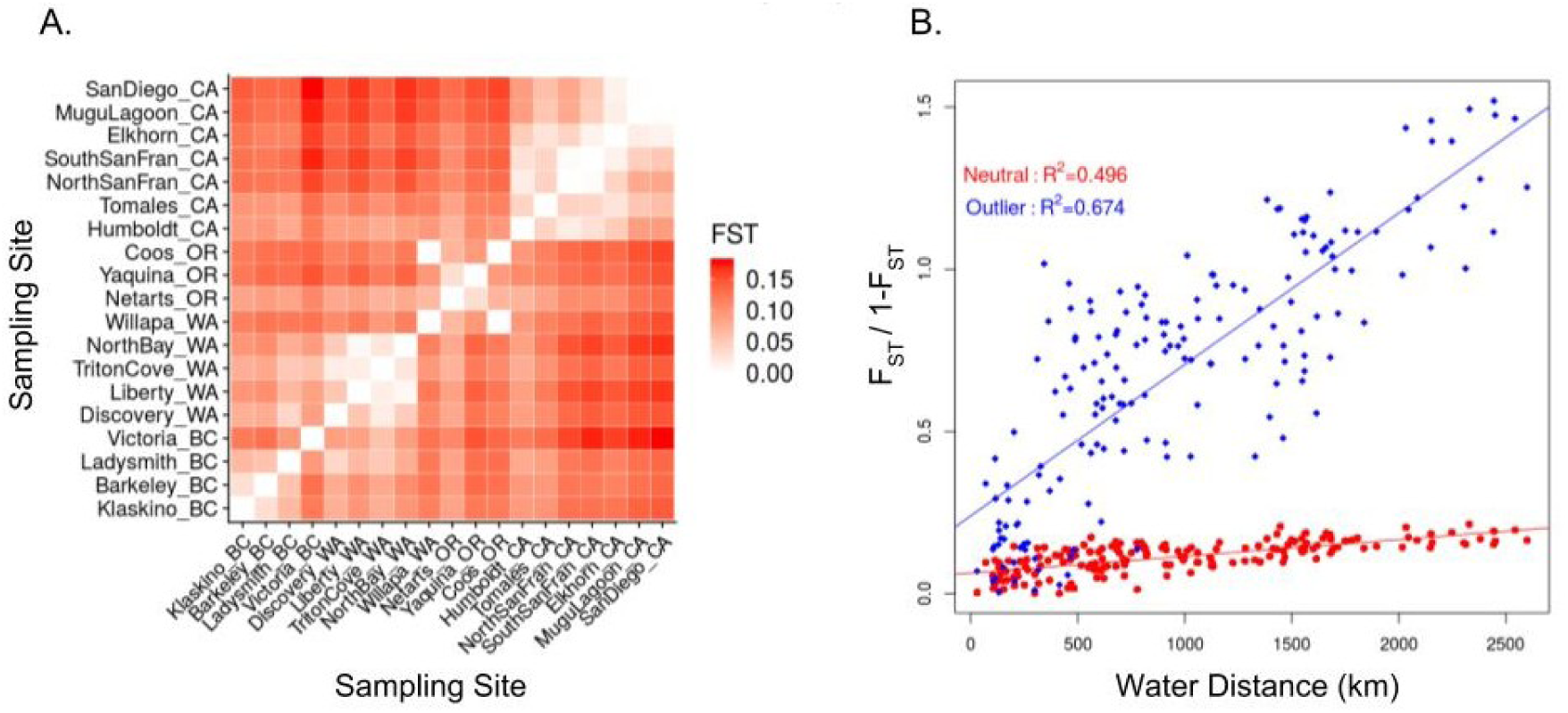
A) Heatmap of pairwise *F*_*ST*_ values for 19 populations of *O. lurida* using 13,093 neutral SNPs. Populations are ordered from north to south, starting with Klaskino, BC. B) Isolation by distance plot of *F*_*ST*_ */*1 *-F*_*ST*_ versus population pairwise coastal water distance. Neutral loci in red circles (*p* < 0.001) and outlier loci in blue diamonds (*p* < 0.001).

### Connectivity and historical relationships

TreeMix produced a population graph that supported a major phylogeographic split between the outer coast of Washington and Puget Sound. When allowing for an increasing number of migration events, the proportion of variance in relatedness between populations explained by the model began to asymptote at 0.991 for 7 migration edges (Figure 4). All of these migration events involve either the *Puget+BC* or *Willapa* regions, except for one from a *NoCal* population to a *SoCal* population. Because the Coos Bay, OR population is likely a recent anthropogenic introduction from Willapa Bay, Coos Bay oysters were excluded from the EEMS analysis. The combined EEMS map for all runs identified four significant (posterior probability > 95%) barriers to gene flow: 1) at the mouth of the Strait of Juan de Fuca, 2) around Victoria, BC, 3) extending from Willapa Bay, WA to southern Oregon, and 4) around San Francisco Bay, CA (Figure 5). These inferred barriers further lend credence to the 6 phylogeographic regions identified through other means. An area of significantly increased gene flow was inferred between Mugu Lagoon and San Diego. The EEMS method also estimated and mapped the genetic diversity parameter *q*, which is an estimate of the expected within-deme coalescent time and is proportional to average heterozygosity (*H*_*e*_). Populations from Oregon northwards had much lower genetic diversity than those in California. A linear regression of population-specific *H*_*e*_ and latitude using the neutral dataset shows a strong relationship between genetic diversity and latitude (*R*^2^ = 0.86)(Figure 6), as did the outlier dataset (*R*^2^ = 0.871).

**Figure 4:**
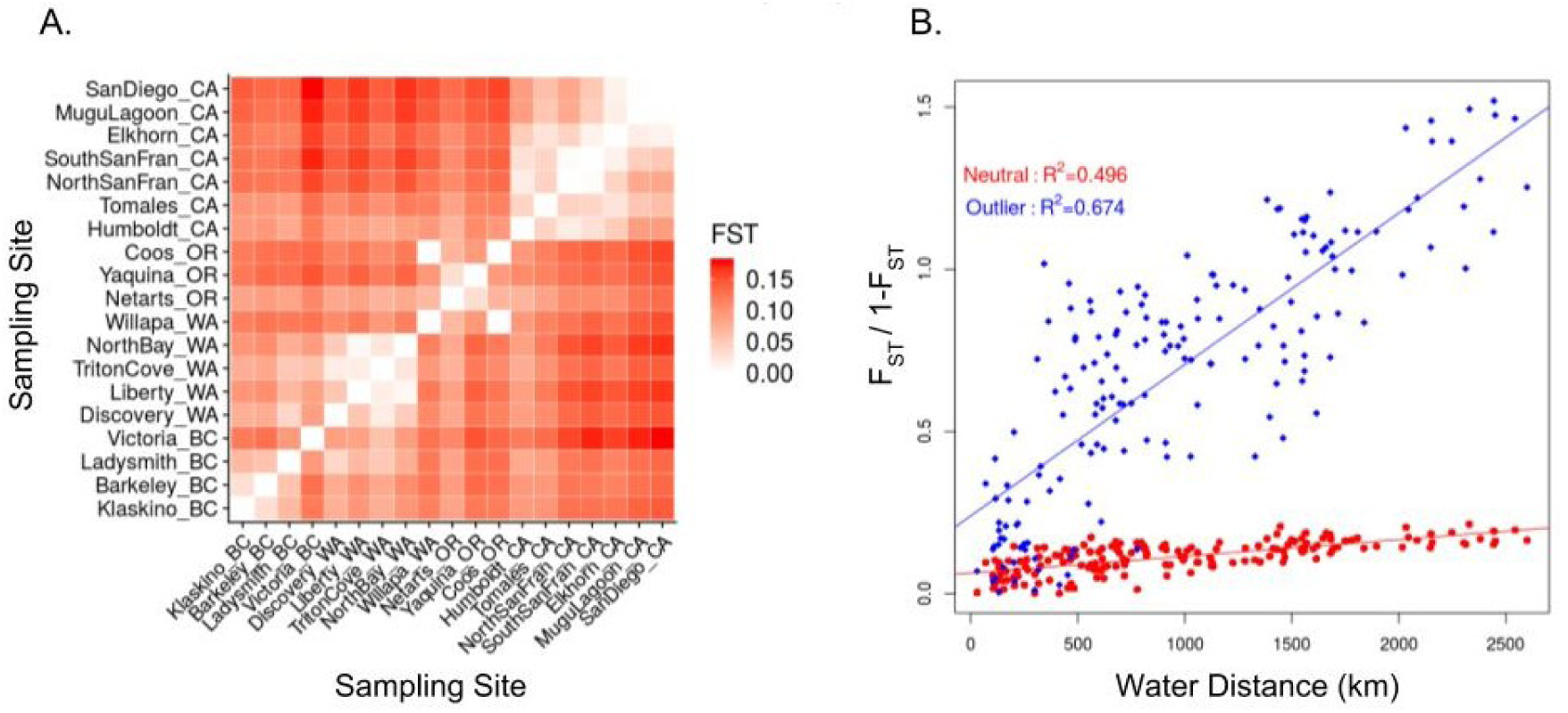
TreeMix results for 19 *O. lurida* populations using 1 SNP per neutral locus. Seven migration events are modeled, as this was the best value inferred by evaluating model fit. The tree is rooted by the southernmost populations, San Diego Bay, CA and Mugu Lagoon, CA, and ordered by latitude where possible. Populations are colored by their inferred phylogeographic region.

**Figure 5:**
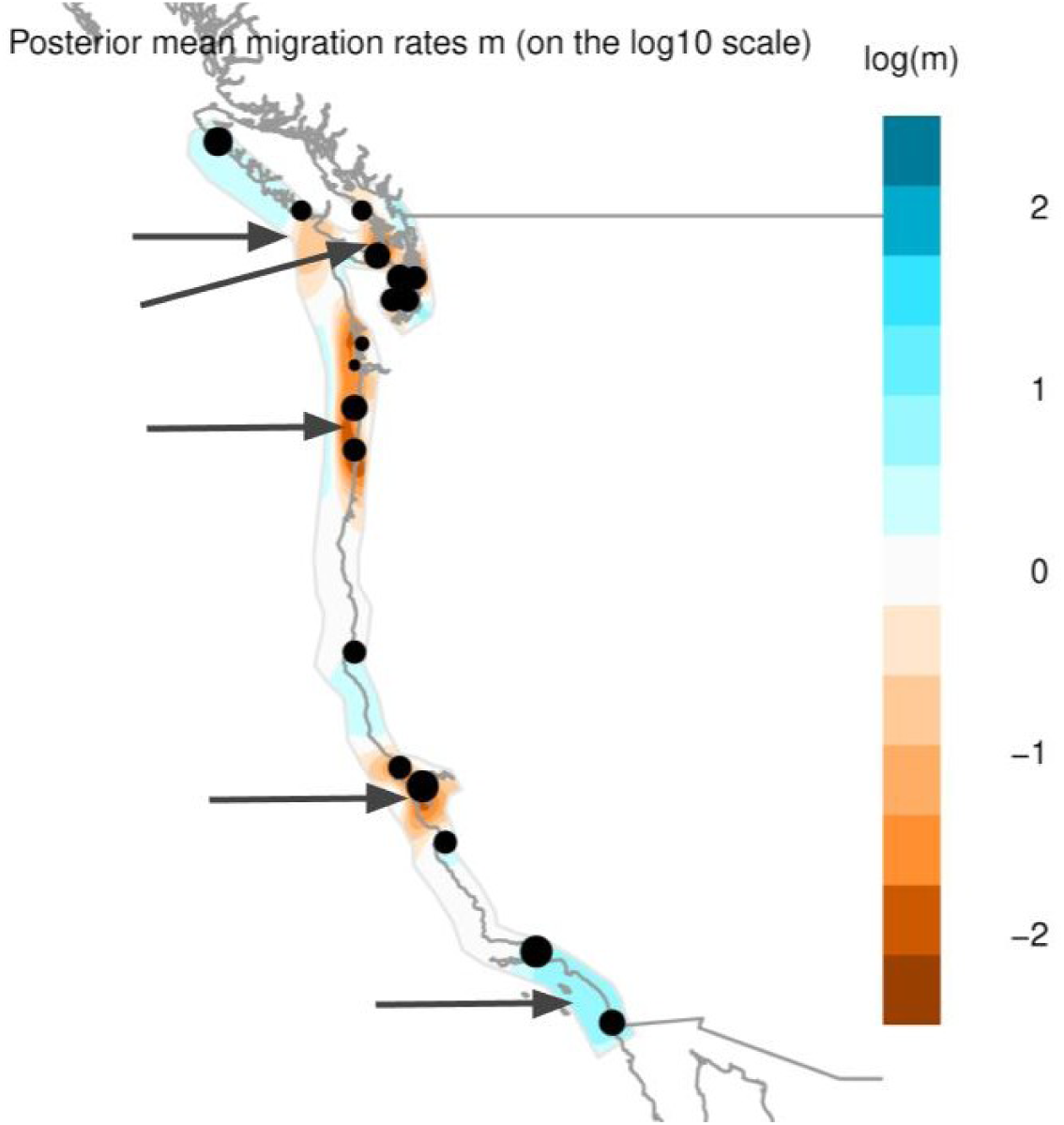
Model of effective migration rates as inferred by EEMS for neutral loci in *O. lurida*. Orange represents ares of low migration relative to the average and blue are areas of higher migration. Grey arrows indicate regions of significantly reduced or increased migration (posterior probability > 95%).

**Figure 6:**
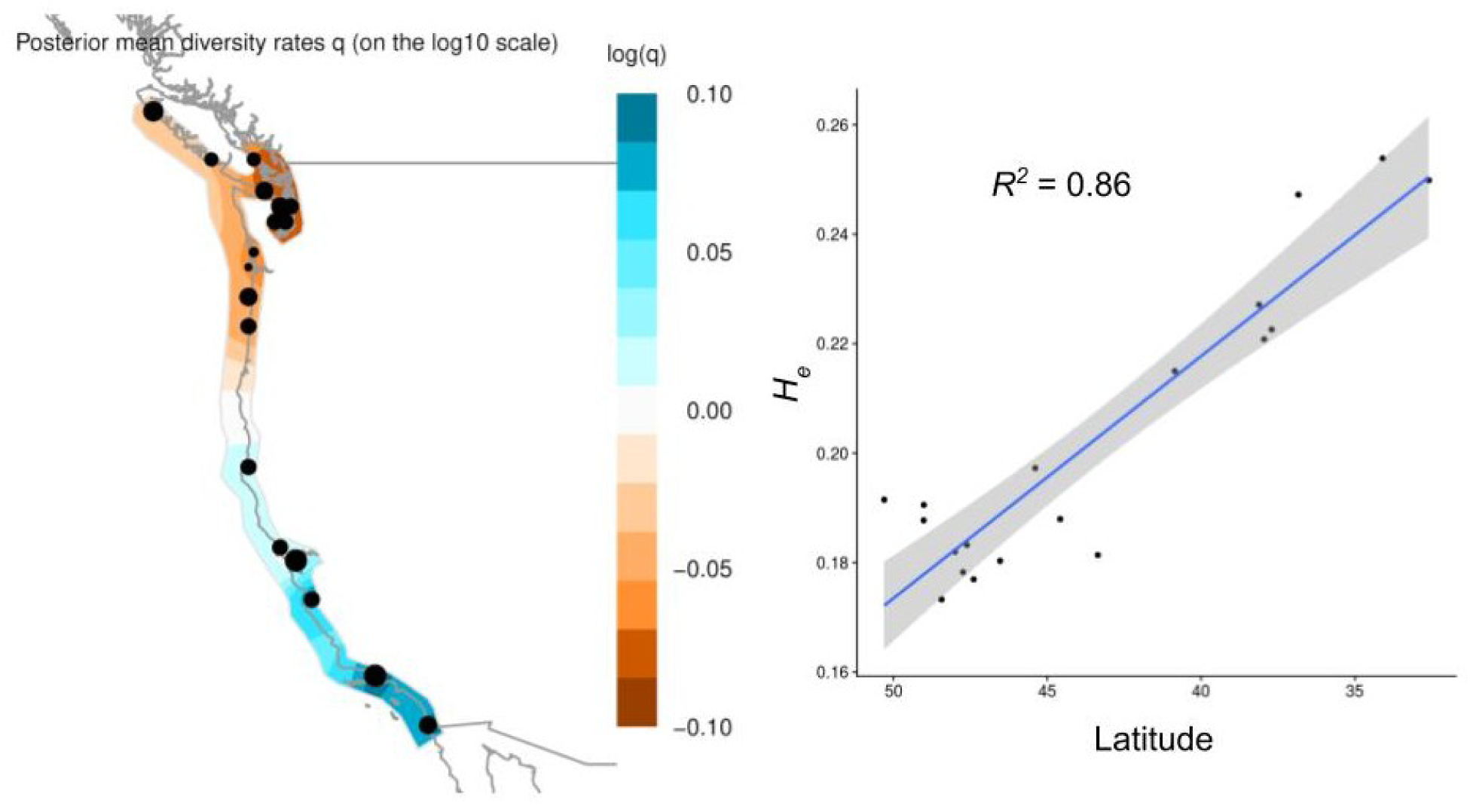
Diversity increases from north to south in *O. lurida*. (A) Effective diversity rates as inferred by EEMS, with orange representing areas of lower diversity and blue representing high diversity. (B) Expected heterozygosity (*H*_*e*_) within each population versus population latitude.

### Functional annotation of outliers

The 129 GBS loci containing outlier SNPs were functionally annotated using Blast2GO. Eighteen of these mapped to protein coding genes in the GenBank database, primarily from *Crassostrea virginica* and *Crassostrea gigas*. One mapped to the *O. lurida* mitochondrial NADH dehydrogenase subunit 5 gene (*nad5*), which exhibits high variability in oyster species and is commonly used for metazoan phylogenetics (Xiao et al., 2015). Annotated genes have potential roles in developmental regulation (glyoxalase 3, DNA N6-methyl adenine demethylase-like, transcriptional regulator ERG, serine/threonine-protein kinase), sensory information processing (serine/threonine-protein kinase, sodium-dependent phosphate transport, vesicular glutamate transporter), immune or stress response (*nad5*, E3 ubiquitin-protein ligase, Ty3-G Gag-Pol, helicase domino), energy metabolism (carnitine palmitoyltransferase, glucose dehydrogenase), heavy metal binding (Heavy metal-binding HIP), and muscle contraction (myosin heavy chain-striated muscle, myosin-XVIIIa) (Table 3) Epelboin et al. (2016); Szent-Gyö rgyi et al. (1999); Li et al. (2017); Cheng et al. (2016); Anderson et al. (2015); Riviere et al. (2013); Pauletto et al. (2017); Wang et al. (2018); Shiel et al. (2017); Pan et al. (2015); de Lorgeril et al. (2005). 21 additional outlier GBS loci had positive matches to InterPro signatures without any BLASTx hits or gene ontology annotation. Plotting minor allele frequency against latitude for outlier SNPs demonstrates that the majority of outliers show a clinal pattern, where one allele is fixed from either Coos Bay, OR or San Francisco Bay, CA to the north, and the other alelle increases in frequency towards the south (Figure D2).

## DISCUSSION

Reduced-representation genomic methods, such as GBS, can greatly inform reintroduction efforts for threatened and exploited species by resolving fine-scaled population structure, providing estimates of genetic connectivity, and identifying informative markers for characterizing adaptive variation (Allendorf et al., 2010; Gagnaire et al., 2015). Using 13,444 GBS-derived SNPs, I characterized the rangewide population structure of the Olympia oyster from southern California to British Columbia and further identified 288 SNPs across 129 GBS loci potentially associated with local adaptation. Contrary to studies in some other marine species, neutral markers had greater power to detect fine-scale population structure compared to outliers. However, outlier loci did provide evidence for adaptive divergence among some populations with high inferred admixture, and are informative as candidate loci involved in local adaptation. This study highlights the importance of using both neutral and outlier markers for conservation and management applications.

### Regional population structure and gene flow

Significant population structure was observed across the range of *O. lurida* in both the neutral and outlier markers, with sampling locations structured into six distinct regions. Notably, most of these regions fit well within previously described biogeographical provinces based on marine species distributions(Hall, 1964; Valentine, 1966; Fenberg et al., 2015). In addition to describing the rangewide population structure of *O. lurida*, the large geographic sampling of this study can facilitate the identification of oceanographic features along the eastern Pacific coast that may be important for structuring populations of marine species with similar life histories. Most of the inferred phylogeographic regions are bounded by areas of reduced gene flow, many of which align to oceanographic features that may be acting as barriers to dispersal. Below I discuss these phylogeographic regions and potential barriers in more detail, as well as provide some recommendations for management at local scales.

### Southern California (SoCal)

The *SoCal* region, containing San Diego Bay, CA, Mugu Lagoon, CA, and Elkhorn Slough, CA, extends across both the Southern Californian and the Montereyan biogeographic provinces as defined by Hall (1964), with Monterey Bay as the northern boundary. Monterey Bay is a known biogeographic barrier for some marine algae (Abbott and Hollenberg, 1976), and has been proposed as a potentially important barrier to gene flow in marine invertebrates as well (Dawson, 2001). This region extends across Point Conception, which is a well-known site of species turnover (Valentine, 1966) and a phylogeographic barrier for some taxa (Marko, 1998; Wares et al., 2001). This finding is consistent with meta-analyses demonstrating that strong population structure across Point Conception is the exception rather than the rule for many marine invertebrates(Kelly and Palumbi, 2010; Dawson, 2001) Finer-scaled sampling of populations on either side of Point Conception may provide evidence for slight genetic clines undetected by the current study.

*SoCal* exhibits the highest genetic diversity of any region, for which I propose three nonexclusive mechanisms. 1) The southward direction of the California Current results in asymmetric gene flow and an accumulation of genotypes in the south (Wares et al., 2001). This hypothesis is supported by the inferred directionality of migration events in TreeMix (Figure 4). 2) Northern populations exhibit lower genetic diversity due to repeated extirpation or population bottlenecks from glaciation cycles (see *Puget+BC*) (Marko, 2004). 3) Ongoing or historical admixture from the southern sister species *O. conchaphila* has increased genetic diversity in these populations. Sampling and genotyping of *O. conchaphila* is underway to test this hypothesis. The low *F*_*ST*_ between Mugu Lagoon and San Diego suggests either a recent transplantation between sites or high gene flow. If the former, I hypothesize that Mugu Lagoon is the recent transplant due to a high inbreeding coefficient (*F*_*IS*_). Nevertheless, three outlier loci exhibited allele frequency shifts of at least 50% between these two populations, suggesting some potential adaptive population divergence.

### Northern California (NoCal)

San Francisco Bay, Tomales Bay, and Humboldt Bay constitute the *NoCal* region, which is encompassed by the northern half of the Montereyan biogeographic province as identified by (Fenberg et al., 2015) and delineated by Cape Mendocino to the north. Cape Mendocino, located 46 km south of Humboldt Bay, is an established phylogeographic break for multiple marine species (Kelly and Palumbi, 2010). EEMS identifies an area of significantly reduced gene flow surrounding San Francisco Bay, which may correspond to Monterey Bay (Dawson, 2001), or anthropogenic introductions (see *Anthropogenic influences on population structure*). The two sites within San Francisco Bay (Candlestick Park and Point Orient), exhibit potential adaptive divergence at some outlier GBS loci despite high potential for gene flow. This result supports evidence for local adaptation from reciprocal transplant studies within San Francisco Bay (Bible and Sanford, 2016), and highlights the importance of taking individual pairwise *F*_*ST*_ values (Tables C1, C2) into account when making reintroduction decisions. TreeMix inferred significant migration between San Francisco Bay and Elkhorn Slough, however migration is likely not consistent between these populations based on synchrony of recruitment dynamics(Wasson et al., 2016).

### Oregon and Willapa

Both the *Oregon* region, comprised of Netarts Bay and Yaquina Bay, and the *Willapa* region with Willapa Bay, WA and Coos Bay, OR, fall within the Mendocinian biogeographic province, which is usually demarcated by either Cape Flattery (Blanchette et al., 2008) or Vancouver Island (Fenberg et al., 2015) to the north. Evidence from TreeMix and Structure indicate that these two regions have a shared phylogeographic history—likely a combination of evolutionary and anthropogenic processes. EEMS robustly infers an area of significantly reduced migration from Willapa Bay, WA to southern Oregon, which I hypothesize is partly due to the high retention of oyster larvae within Willapa Bay during the summer reproductive season (Banas et al., 2009). EEMS also infers an area of slightly increased migration to the west of the sampling sites. This result may be the EEMS model attempting to incorporate evidence for long-range migration events, likely anthropogenic in nature, between *Puget+BC* sites and sites in *Oregon* and *NoCal*, or it may be an artifact of the model. To my knowledge, this is the first application of EEMS to a marine invertebrate—simulations and additional empirical studies are necessary to evaluate the behavior of EEMS in linear habitats. Currently the protections against importing shellfish from outside of the state are higher than moving shellfish within the state. The strong phylogeographic divide between Willapa Bay, WA and Puget Sound, WA presented here indicates that transfer of Olympia oysters or *Crassostrea* shells between the outer coast of WA and Puget Sound should be considered equivalent to importing oysters from out of state.

### Puget Sound, WA and British Columbia (Puget+BC and NWBC)

The *NWBC* region, comprised of Klaskino Inlet, BC and Barkeley Sound, BC, is significantly differentiated from other sites on Vancouver Island and shows evidence for decreased migration out of the region. The *Puget+BC* region is comprised of Ladysmith Harbour, BC, Victoria Gorge, BC, and all four sites in Puget Sound, WA. Strong evidence suggests that Victoria Gorge, BC has a shared evolutionary history with Puget Sound, WA, although EEMS indicates that migration is reduced between these sites. Ladysmith Harbour may belong to a separate phylogeographic region all together, as this site was intermediate between *NWBC* and *Puget+BC* regions in the STRUCTURE, PCA, and TreeMix analyses. Genetic sampling from additional sites on the central coast of British Columbia and eastern coast of Vancouver Island could test this hypothesis.

The separation of these two regions from those to the south corroborates previous evidence from mitochondrial loci of a strong phylogeographic divide (Polson et al., 2009). Although Cape Flattery and Puget Sound itself have both been classified as biogeographic barriers due to a bifurcation in ocean currents (Valentine, 1966; Kelly and Palumbi, 2010), there are surprisingly few studies evaluating the genetic structure of species found both within Puget Sound and on the outer coast of Washington. Those that do focus on species with much longer dispersal times than *O. lurida* (Buonaccorsi et al., 2002; Cunningham et al., 2009; Iwamoto et al., 2015; Siegle et al., 2013; Jackson and O’Malley, 2017). To my knowledge, this is the first study in a marine mollusc to evaluate and identify significant population differentiation among Puget Sound populations and the outer coast. More studies are required to fully characterize the importance of this barrier across marine taxa. Genetic differentiation within Puget Sound is relatively low at both neutral and outlier markers, with the exception of the northernmost site, Discovery Bay. The weak population structure within Puget Sound and the overall low genetic diversity in northern sites is likely due to recent genetic bottlenecks and range expansion after the last glacial maximum, which reached just north of Willipa Bay, WA (49°N latitude) until 12-13 kya (Dyke and Prest, 1987). Despite such low genetic differentiation, experimental assessments of local adaptation for populations within Puget Sound have detected heritable differences in fitness traits such as reproductive timing, growth rate, and gene expression in response to stress (Heare et al., 2017, 2018; Silliman et al., 2018). These results, coupled with experimental evidence for local adaptation to salinity among Northern California populations (Bible and Sanford, 2016), suggest that adaptive divergence in this species can occur in the face of high gene flow.

### Anthropogenic influences on population structure

The evidence for reduced effective migration, low differentiation within most of the phylogeographic regions, and external estimates of effective dispersal (Carson, 2010), suggests that long distance dispersal is not a significant force in shaping population structure in this species. However, TreeMix inferred a few such migration events that cross aforementioned barriers to gene flow. To explain this evidence, I investigated the history of Olympia oyster exploitation and aquaculture through literature reviews, technical reports, grey literature, historical first-person accounts, and discussions with current restoration practitioners. The historical impact of human take and transportation on the Olympia oyster is substantial.

Beginning in 1850, oysters were shipped from Willapa Bay to northern California by the millions, including shipments of juvenile or “seed” oysters to be raised in local waters until reaching commercial size. The inferred migration events from North Bay, WA to California sites may be reflecting the historical transplantation of seed oysters from Oakland Bay, WA, about 30 km from North Bay(Woelke, 1959; Baker, 1995). After the crash of the Olympia oyster industry, the non-native oysters *Crassostrea virginica* and *C. gigas* were brought to the west coast for commercial aquaculture. The shells of these species are excellent substrate for Olympia oysters, and the movement of *Crassostrea* oysters between bays for culturing purposes (e.g., San Francisco to Humboldt in 1910, Willapa to Humboldt in 1950s (Barrett, 1963)) may have resulted in the accidental transfer of *O. lurida* (Townsend, 1895). The low *F*_*ST*_ of Willapa Bay, WA and Coos Bay, OR, despite being separated by 415 km, corroborates the theory of an accidental introduction of Olympia oysters from Willapa Bay on *C. gigas* shells in the 1980s (Baker et al., 1999). The low inbreeding coefficient for Coos Bay suggests a potentially large founding population. Future movement of *Crassostrea* for aquaculture purposes should be carefully monitored to prevent the accidental migration of nonnative *O. lurida* genotypes.

While it is encouraging that programs such as TreeMix can recover known human-mediated migration events, such artificial movement of individuals can complicate the determination of natural connectivity patterns. For example, the area of low effective migration inferred around San Francisco Bay may be due to the introgression of Washington genotypes rather than actual physical barriers to gene flow. Fortunately, intentional movement of Olympia oysters between regions ceased over 80 years ago with the exception of restoration efforts in Netarts Bay from 2005-2012 which utilized broodstock from Willapa Bay.

### Local adaptation

Detection of outlier loci using three different methods conservatively identified 129 GBS loci as under putative selection. Only 18 GBS loci mapped to protein-coding regions, 3 of which were identified as outliers by all three approaches. Mapping of outlier GBS loci to the forthcoming *O. lurida* genome will aid in detecting loci that may be tightly linked to a gene or regulatory region. Direct information about the function of genes or proteins in oysters is sparse, but rapidly increasing with transcriptomic and physiological studies on the commercially important *Crassostrea* species. While these 129 GBS loci are likely only a fraction of all loci under divergent selection across the *O. lurida* genome (Lowry et al., 2017) and their functional associations are strictly hypotheses, they are nonetheless excellent candidates for future directed studies.

Plotting minor allele frequency against latitude demonstrates that the majority of outlier SNPs show a clinal pattern, where one allele is fixed north of Coos Bay, OR and the other increases in frequency towards the south (Figure D2). Many clinal GBS loci with functional annotations are associated either directly or indirectly with development. Glyoxalase 3 expression has been linked to developmental competence in female oyster gametes (Pauletto et al., 2017), and DNA N6-methyl adenine demethylase has been linked to developmental timing in oysters (Riviere et al., 2013). ERG transcriptional regulator, kinase B-raf, and Fez family zinc finger protein are likely to be directly involved with developmental regulation (Gaitán-Espitia and Hofmann, 2017; Epelboin et al., 2016). Olympia oysters exhibit latitudinal variation in gonad development and spawning, with California oysters initiating spawning up to 6°C warmer than those from Puget Sound, WA (Coe, 1932; Hopkins, 1937). Recent evidence suggests that there is heritable, adaptive variation in reproductive timing, even among populations of oysters within the same phylogeographic region (Silliman et al., 2018; Barber et al., 2016). Another clinal locus of interest mapped to the mitochondrial gene carnitine Ol-palmitoyltransferase, which has been strongly associated with regulation of glycogen content (and therefore, tastiness) in *C. gigas* (Li et al., 2017). Other clinal genes have putative functions in sensory information processing and muscle contraction.

Some outlier loci exhibit the opposite of a clinal pattern, where populations in the middle of the range predominantly have a different allele than the northern and southern populations. E3 ubiquitin-protein ligase (locus 121489) is diverged in *Oregon* and Willapa Bay, WA compared to the other populations. An E3 uniquitin-protein ligase was recently identified as an important component of the neuroendicrine-immune response in *C. gigas* and is primarily expressed in the gonads. Heavy metal-binding protein (HIP) also exhibits this hump-shaped distribution of allele frequencies.

### Potential Limitations

Although genomic methods such as GBS have proven useful for evolutionary biology and conservation genetic studies (Andrews et al., 2016), several potential limitations of GBS and the present study should be addressed. Non-random missing data due to polymorphisms in the restriction enzyme cut site (”allelic dropout”) can bias population genetic analyses by underestimating genomic diversity and overestimating *F*_*ST*_, however the impact of these biases on *F*_*ST*_ may be limited if effective population size (*N*_*e*_) is small and if loci with large amounts of missing data are removed from analyses(Gautier et al., 2013; Cariou et al., 2016). Due to large variation in reproductive success every generation, *N*_*e*_ is likely small for *Ostrea* species (Hedgecock, 1994; Lallias et al., 2010). Loci with > 25% missing data were removed from population genetic analyses, and preliminary analyses allowing 40%–10% missing data still resulted in the same regional population structure and relative values of pairwise *F*_*ST*_, although absolute values of *F*_*ST*_ changed slightly. Two reasons may underlie the large number of individuals (128) removed during filtering. First, too many individuals may have been pooled per sequencing lane given the number of loci targeted, resulting in low sequencing depth for some individuals (Andrews et al., 2016). Second, these libraries were made and sequenced in-house as opposed to a dedicated commercial GBS facility. The protocol learning curve may be why a disproportionate number of individuals failed or had low sequencing depth in the first few prepared libraries. This filtering resulted in 4–9 individuals per population in the final dataset, which is sufficient for estimating *F*_*ST*_ when > 1,000 SNPs are used (Willing et al., 2012). While these small population sizes may limit the power to detect outlier loci (Foll and Gaggiotti, 2008), the probability of false positives is reduced by comparing across multiple outlier methods (Rellstab et al., 2015). Lastly, while methods like EEMS and PCA can characterize genetic differentiation, they cannot distinguish between the different demographic scenarios that may result in these patterns(Petkova et al., 2016).

## CONCLUSIONS

This study provides the first comprehensive characterization of both neutral and adaptive population structure in the Olympia oyster, an ecologically important coastal species in North America. These results have direct implications for management policies and ongoing restoration efforts, and a future sustainable fishery. Putative adaptive loci identified here are excellent candidates for future research and may provide targets for genetic monitoring programs. Beyond these specific applications, this study contributes to the growing body of evidence for both population structure and adaptive differentiation in marine species. In particular, it is one of the first to utilize thousands of SNPs to characterize population structure from southern California to Vancouver Island. All analyses conducted for this study can be replicated using annotated Jupyter notebooks, allowing for clear dissemination of bioinformatics methods and future open-sourced research on the population structure of *O. lurida*.

## DATA ARCHIVING STATEMENT

Reproducible Jupyter notebooks will be available at [redacted]. Raw sequencing data will be stored on the NCBI Sequence Read Archive. Sequence data after all filtering and assembly will be available on Dryad.

## ACKNOWLEDGEMENTS

I would like to thank the Washington Dept. of Fish and Wildlife, Jennifer Ruesink, Alan Trimble, Danielle Zacherl, Thomas McCormick, JoAnne Linnenbrink, Daniel Silliman, Brian Kingzett, Steven Rumrill, and Dave Couch for help and advice with sample collections, as well as the Field Museum Pritzker DNA Lab for providing reagents, facilities, and funding for molecular work. Additional funding was provided by the University of Chicago Hinds Fund and student awards through the National Shellfisheries Association and the American Fisheries Society. This material is based upon work supported by the National Science Foundation Graduate Research Fellowship under Grant No. 1545870 and the Department of Education Graduate Assistance in Areas of National Need Fellowship.

## APPENDICES

### Appendix A: Sampling locations and population-specific summary statistics

**Table A.1.**
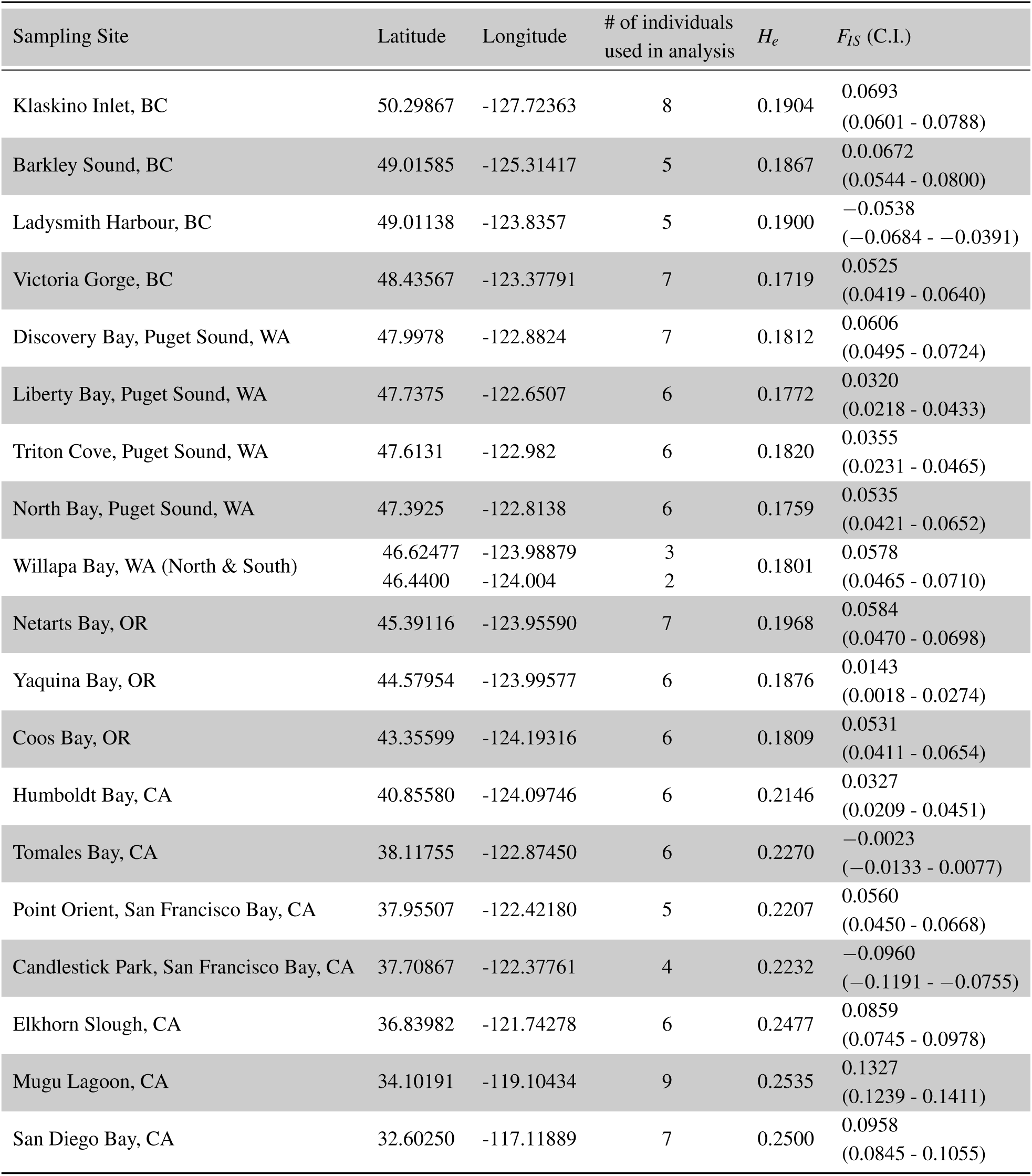
GPS coordinates of sampling sites and population-specific summary statistics averaged across markers using the combined dataset of 13,444 SNPs. *H*_*e*_, expected heterozygosity; *F*_*IS*_, inbreeding coefficient within the population, mean and 25%-75% confidence intervals (Nei and Chesser, 1983);

### Appendix B: Summary statistics for phylogeographic regions

**Table B.1.** Overall summary statistics for each phylogeographic region using the neutral dataset of 13,093 SNPs. *H*_*o*_, observed heterozygosity averaged across loci; *H*_*e*_, expected heterozygosity averaged across loci; *F*_*IS*_ & *F*_*ST*_, Wright’s *F*-statistics averaged across loci (Nei and Chesser, 1983). Note that *F*_*ST*_ may be skewed by variation in sampling strategy across regions.

| Region | $H_o$ | $H_e$ | $F_{IS}$ | $F_{ST}$ |
| --- | --- | --- | --- | --- |
| NWBC | 0.177 | 0.193 | 0.0821<br>(0.0738 - 0.0897) | 0.016 |
| Puget+BC | 0.174 | 0.189 | 0.0814<br>(0.0758 - 0.0862) | 0.046 |
| Willapa | 0.171 | 0.182 | 0.0583<br>(0.0495 - 0.0666) | 0.001 |
| Oregon | 0.185 | 0.196 | 0.0556<br>(0.0474 - 0.0645) | 0.016 |
| NoCal | 0.215 | 0.227 | 0.0536<br>(0.0472 - 0.0592) | 0.022 |
| SoCal | 0.224 | 0.253 | 0.115<br>(0.1097 - 0.1209) | 0.007 |

### Appendix C: Pairwise FST values for outlier and neutral datasets

**Table C.1.**
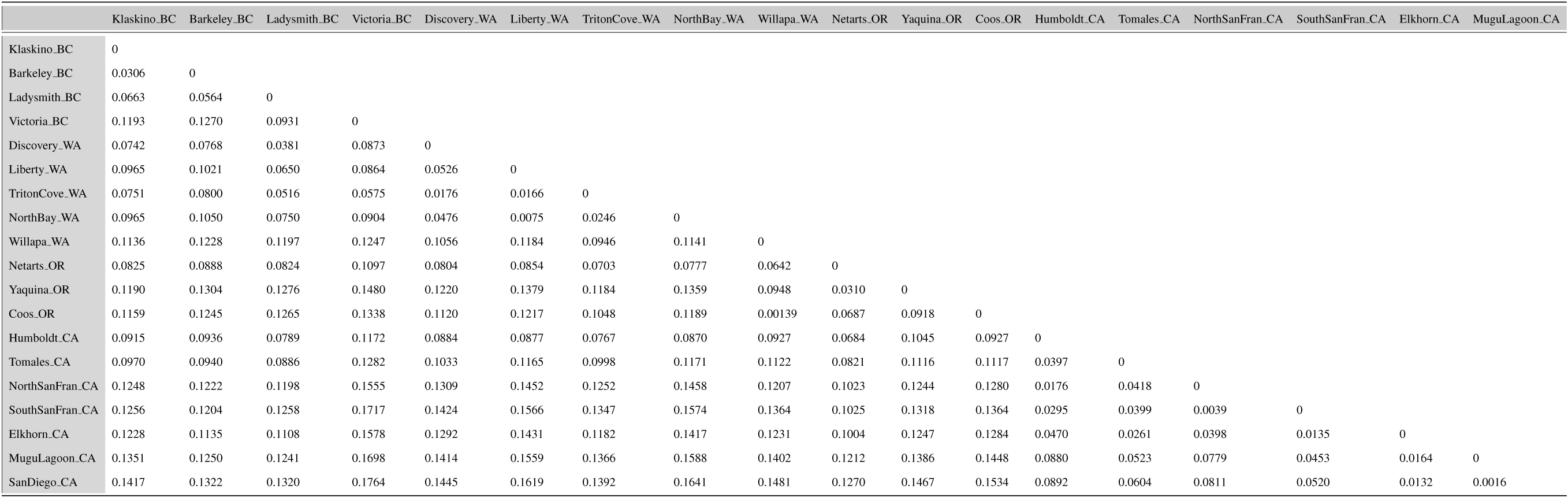
Pairwise *F*_*ST*_ values for all pairs of populations, using the neutral dataset of 13,093 SNPs.

**Table C.2.**
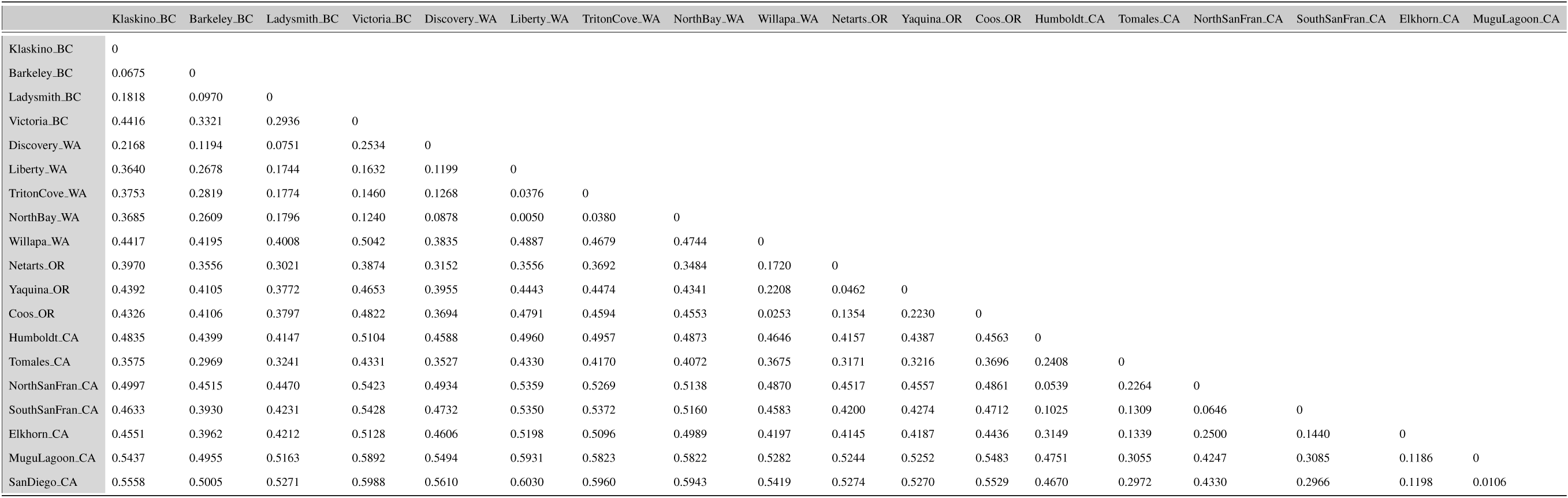
Pairwise *F*_*ST*_ values for all pairs of populations, using the outlier dataset of 288 SNPs.

### Appendix D: Additional results of outlier analyses

**Figure D.1.**
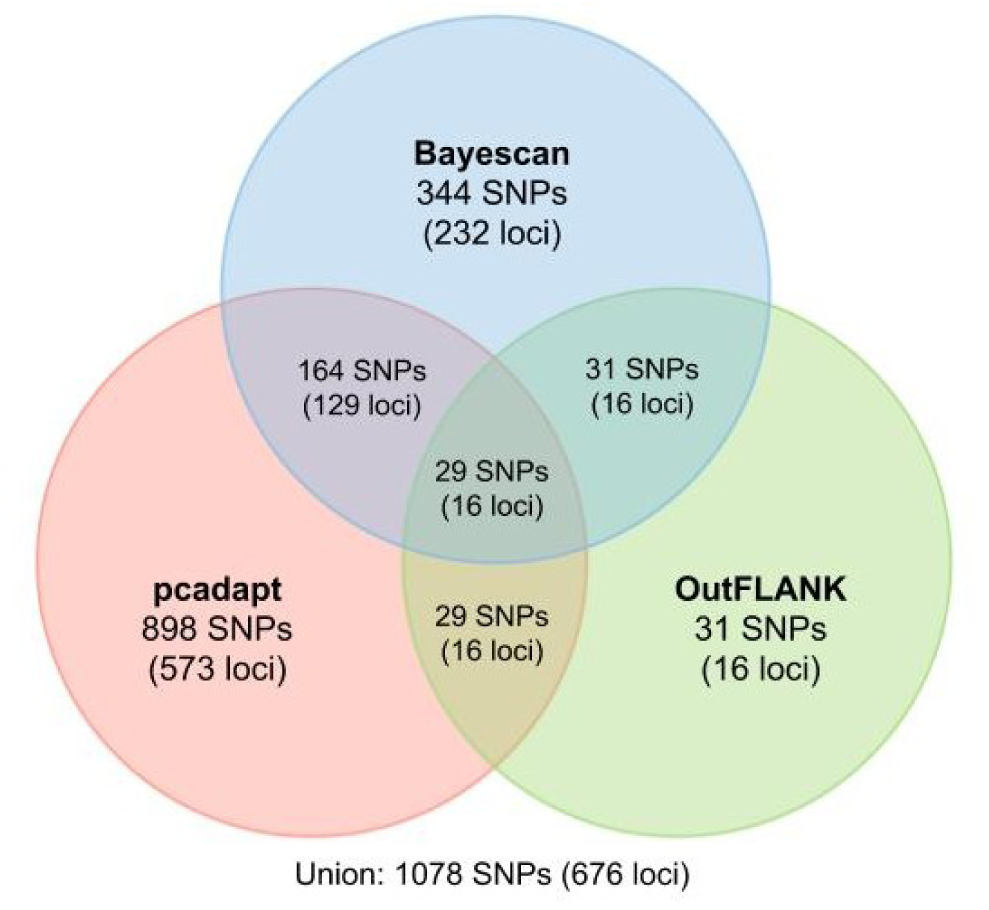
Venn diagram with number of SNPs and Genotype-by-Sequencing loci identified as outliers by three methods: *pcadapt, OutFLANK*, and BayeScan

**Figure D.2.**
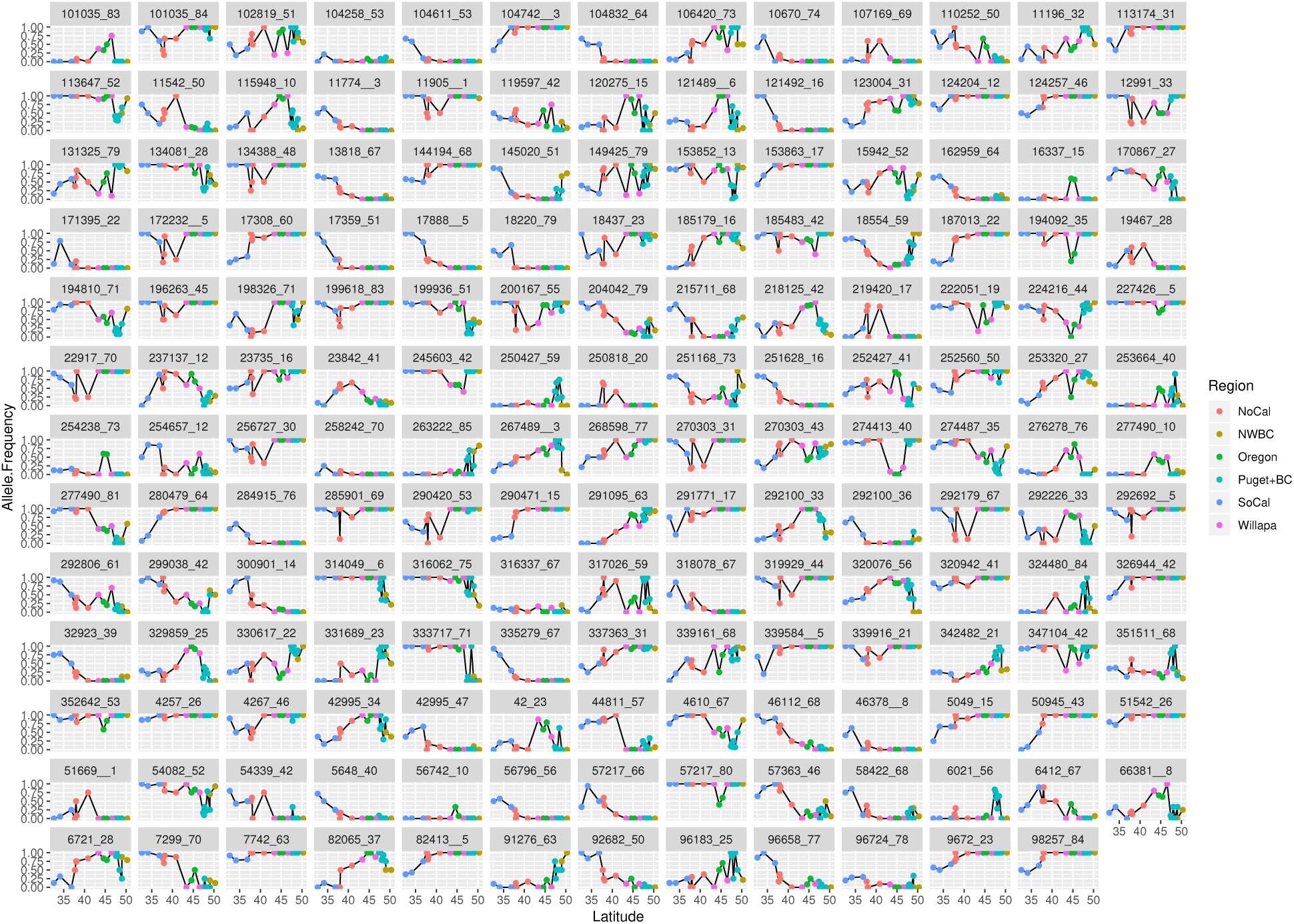
Outlier loci predominantly show clinal patterns in allele frequency. Allele frequency in 129 individual outlier loci plotted against latitude for 19 populations of *O. lurida*. One SNP is represented for each locus, except in the case where two outlier SNPs from the same locus showed different spatial patterns (e.g., locus 277490). Populations are colored by inferred phylogeographic regions.

